# Thalamocortical and Intracortical Laminar Connectivity Determines Sleep Spindle Properties

**DOI:** 10.1101/195552

**Authors:** Giri P Krishnan, Burke Q Rosen, Jen-Yung Chen, Lyle Muller, Terrence J Sejnowski, Sydney S Cash, Eric Halgren, Maxim Bazhenov

## Abstract

Spindle oscillations are brief oscillatory activity during non-rapid eye movement (NREM) sleep. Spindle density and synchronization properties are different in MEG versus EEG recordings in humans and also vary with learning performance, suggesting spindle involvement in memory consolidation. Using computational models, we identified network mechanisms that may explain differences in spindle properties across cortical structures. First, we report that differences in spindle occurrence between MEG and EEG data may arise from the properties of the core vs. matrix thalamocortical systems. The matrix system, projecting superficially, has wider thalamocortical fanout compared to the core system, projecting to the middle layers, and requires the recruitment of a larger population of neurons to initiate a spindle. Our model demonstrates that this property is sufficient to explain lower spindle density and higher spatial synchrony of spindles in the superficial cortical layers, as observed in the EEG signal. In contrast, spindles in the core system occurred more frequently but less synchronously, as observed in the MEG recordings. Futhermore, consistent with human recordings, in the model, spindles occurred independently in the core system but matrix system spindles commonly co-occurred with core one. We found that the intracortical excitatory connections from layer III/IV to layer V promote spindle propagation from the core to the matrix system, leading to widespread spindle activity. Our study predicts that plasticity of the intra and inter cortical connectivity can potentially be a mechanism for increasing in spindle density as observed during learning.

**Author summary:** The density of sleep spindles has been shown to correlate with memory consolidation. Further, sleep spindles occur more often in human MEG than EEG. We developed thalamocortical network model that is capable of spontaneous generation of spindles across cortical layers and that captures the essential statistical features of spindles observed in experiments. We predict that differences in thalamo-cortical connectivity, known from anatomical studies, lead to the differences in the spindle properties between EEG and MEG as observed in human recordings. Further, we predict that the intracortical connectivity between cortical layers, a property influenced by sleep preceding learning, increases spindle density. Results from our study highlight the role of cortical and thalamic projections on the occurrence and properties of spindles.

## Introduction

Sleep marks a profound change of brain state as manifested by the spontaneous emergence of characteristic oscillatory activities. In humans, sleep spindles consist of waxing-and-waning bursts of field potentials oscillating at 11–15 Hz lasting for 0.5–3 s and recur every 5–15 s. Experimental and computational studies have identified that both the thalamus and the cortex are involved in the generation and propagation of spindles. Spindles are known to occur in the isolated thalamus after decortication *in vivo* and thalamic slices recordings in vitro [1, 2], implicating the thalamus as a minimal spindle generator. In *in-vivo* conditions, the cortex has been shown to be actively involved in the initiation and termination of spindles [3] and long-range synchronization of spindles [4] [5].

Multiple lines of evidence indicate that spindle oscillations are linked to memory consolidation during sleep. Spindle density is known to increase following training in hippocampal dependent memory task [6] and during procedural memory [7]. The spindle density is also shown to correlate with better memory retention following sleep in verbal tasks [8, 9]. More recently, it was shown that pharmacologically increasing spindle density leads to better post-sleep performance in hippocampal-dependent learning tasks [10]. In contrast, these experiments found no relationship between spindle amplitude and learning performance, suggesting spindle event occurrence (as oppose to spindle amplitude) in memory consolidation.

In human recordings, spindle occurrence and synchronization vary based on the recording modality and spatial location. In MEG, spindles occur more often and are less synchronized, while EEG spindles are less frequent but more synchronized [11]. It has been proposed that the contrast between observed MEG and EEG spindles reflect differences in synchronization across cortical layers [5] resulting from the involvement of the two primary thalamocortical systems [12]. This hypothesis is supported by human laminar microelectrode data in which spatially coherent spindles are observed in upper layers and less coherent spindles occur in the middle layers [13]. Taken together, these studies suggest that there could be two systems of spindle generation within the cortex and that these correspond to the core and matrix anatomical networks. However, how interaction between the two systems lead to both independent and co-occurring spindles across cortical layers is not understood.

In this study, we developed a computational model that replicates known features of spindle occurrence in MEG and EEG human recordings. While our previous efforts have been focused on the neural mechanisms involved in the generation of isolated spindles[5], in this study we identified the critical mechanisms underlying the spontaneous generation of spindles across different cortical layers and their interaction.

## Results

### Spindle occurrence is different in EEG and MEG

Histograms of EEG and MEG gradiometer inter-spindle intervals are shown in Fig. 1C. For neither channel type are ISIs distributed normally as determined by Lilliefors tests (D_2571_ = 0.1062, p =1.0e-3, D_4802_ = 0.1022, p =1.0e-3), suggesting that traditional descriptive statistics are of limited utility. However, the ISI at peak of the respective distributions is longer for EEG than it is the MEG. In addition, a two-sample Kolmogorov-Smirnov test confirms that EEG and MEG ISIs are not drawn from the same distribution (D_2571,4802_ = 0.079, p = 1.5e-9). While the data from where not found to be drawn from any parametric distribution with 95% confidence, an exponential fit (MEG) and lognormal fit (EEG) are shown in red overlay for illustrative purposes. These data are consistent with previous empirical recordings [14] and suggest that sleep spindles have different properties across superficial vs. deep cortical layers.

**Figure 1:**
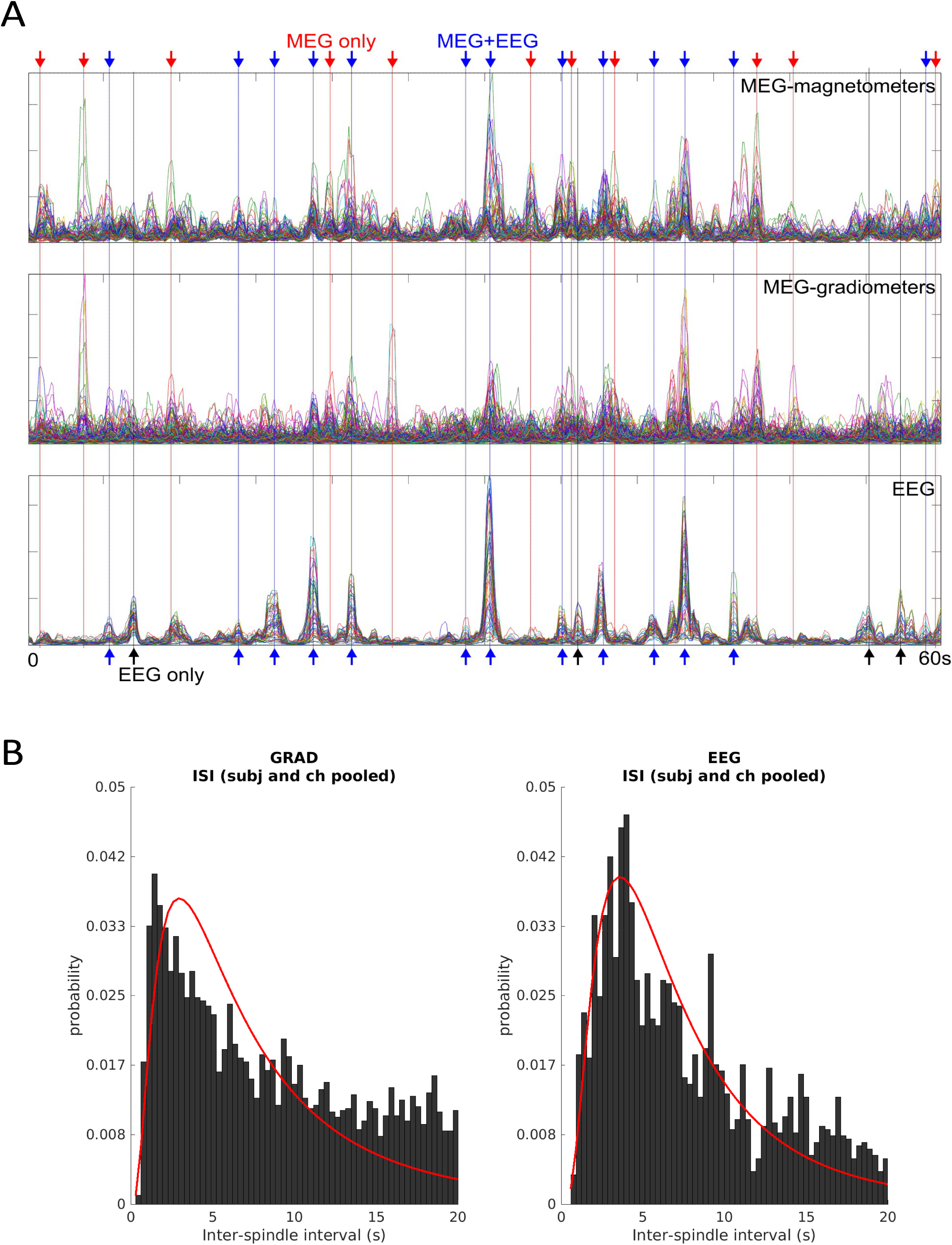
Experimental Observations. (A) Envelopes of spindle-band (6-15Hz) activity in human MEG magnetometers, MEG gradiometers, and EEG. Within-modality channels are superimposed. Arrows indicate spindles that occur in MEG only, EEG only, and both MEG & EEG, in red, black, and blue, respectively. Note that EEG spindles are less frequent and have greater synchrony than MEG spindles. (B) Histograms of the inter-spindle intervals during stage 2 sleep show a non-normal distribution in the occurrence of spindles in both the MEG gradiometer and EEG channels. Compare with simulated data in Fig 4B.

### Frequent local spindle in core and rare global spindles in matrix

To investigate the mechanisms behind distinct spindle properties across the cortical locations as observed in the EEG and MEG signals, we constructed a thalamocortical network model that incorporated two characteristic thalamocortical systems: core and matrix. The two systems contained distinct thalamic populations that projected to the superficial (matrix) and middle (core) cortical layers. Four cell types were used to model distinct cell populations: thalamocortical relay (TC) and reticular (RE) neurons in thalamus, and each layer of cortical network consisted of excitatory pyramidal (PY) and inhibitory (IN) neurons. A schematic representation of the synaptic connections and cortical geometry of our thalamocortical network model is shown in Fig 2. In the matrix system, both thalamocortical (from matrix TCs to the apical dendrites of layer 5 pyramidal neurons (PYs) located in the layer 1) and corticothalamic projections (from layer 5 PYs back to the thalamus) formed diffuse connections. The core system had focal connection pattern in both thalamocortical (from core TCs to the PYs in the layer III/IV) and corticothalamic (from layer VI PYs to the thalamus) projections. Since the spindles recorded in the EEG signal reflect the activity of superficial layers while MEG records spindles originating from the deeper layers (Fig 1 and [13]), we compared activity of the model matrix system, which has projections to the superficial layers to the empirical EEG recordings and compared the activity in model layer 3/4 to the empirical MEG recordings.

**Figure 2:**
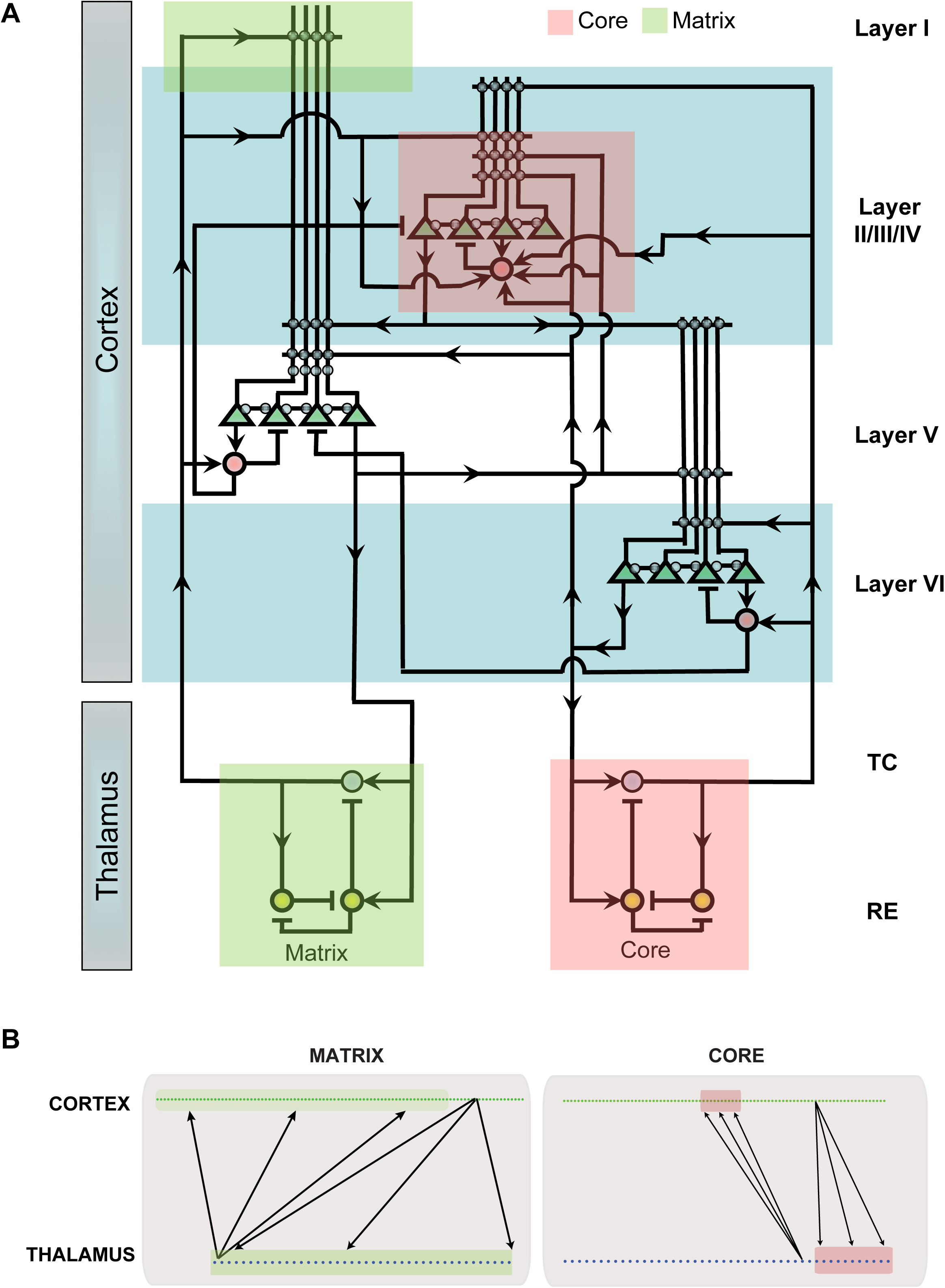
Connectivity in the thalamocortical network. A. The network architecture included three layers of the one-dimensional cortical neurons and two thalamic neuron groups corresponding to the matrix and core systems. Green triangles indicate cortical pyramidal neurons, red circles indicate inhibitory neurons in cortex, blue circles indicate thalamic relay cell and yellow circles indicate thalamic reticular cells. Lines ending with arrows indicate AMPA connections while lines ending with bars indicate inhibitory GABAergic connections. B. Schematic connectivity for the differences in spatial extent of thalamocortical and corticothalamic projections for core and matrix systems.

In agreement with our previous studies [3, 5, 15, 16], simulated stage 2 sleep consisted of multiple spindle events involving thalamic and cortical neuronal populations (Fig 3). During one such typical spindle event (highlighted by the box in Fig 3A and B), cortical and thalamic neurons in both the core and matrix system had elevated and synchronized firing (Fig 3A bottom), consistent with previous in-vivo experimental recordings [17]. In our model, spindles within each system were initiated from spontaneous activity within cortical layers and spread to thalamic neurons similar to our previous study[5]. The spontaneous activity due to miniature EPSPs in glutamergic cortical synapses lead to fluctuations in membrane voltage and sparse firing. At random times, the miniature EPSPs summed such that a small number of locally connected PY neurons spiked within a short window (<100ms), which then induced spiking in thalamic cells through the cortico-thalamic connections. This initiated spindle oscillations in the thalamic population mediated by the TC-RE interaction [15, 18, 19]. Thalamic spindles in turn propagated to the neocortex leading to joint thalamo-cortical spindle events whose features were shaped by the properties of thalamocortical and corticothalamic connections. In this study, we examined how the process of spindle generation occurs in a thalamocortical network with mutually interacting core and matrix systems, wherein the thalamic network within each system is capable of generating spindles independently.

**Figure 3:**
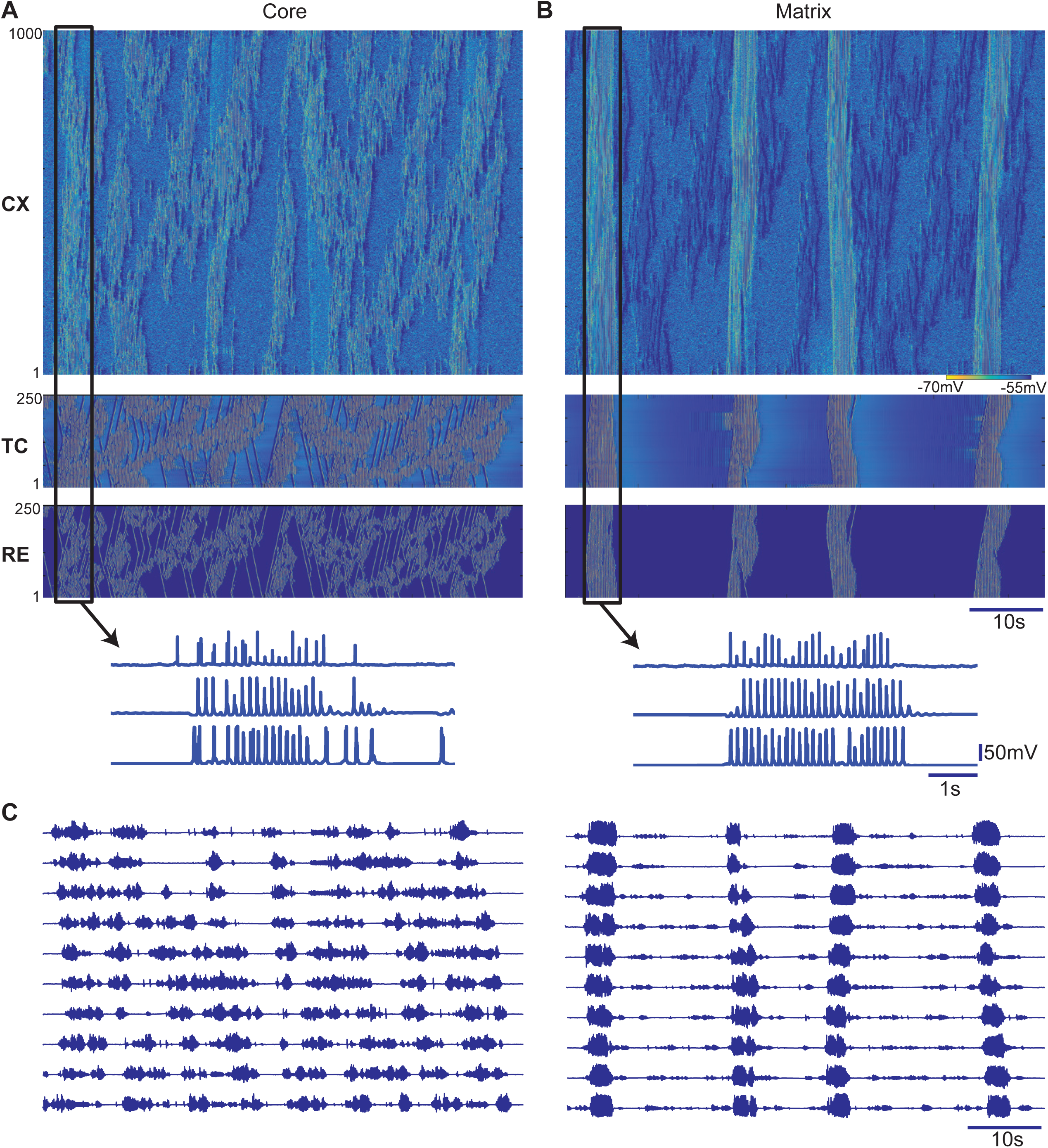
Spindle activity is different in the core and matrix systems. (A, B) Space-time plots of the network activity in this thalamocortical network including core system (A) and matrix system (B). (C) Average network activity (simulated LFP) filtered between 6-15 Hz was measured from 10 non-overlapping locations (each group contains 100 PYs) of the cortical network. Note, spindles occur more frequently (but less synchronously) in L3/4 and less frequent (but more synchronous) in L5.

Based on the anatomical data [12], the main difference between the modeled core and matrix systems was the radii or fanout of connections in thalamocortical and corticothalamic projections (in baseline model, the fanout was 10 times wider for matrix compared to the core system). Furthermore, the strength of each synaptic connection was scaled by the number of input connections to each neuron [20, 21], leading to more weakly connected thalamocortical projections in the matrix as compared to the core. These differences in strength and fanout of the thalamus-cortex connectivity resulted in distinctive core and matrix spindle properties (see Fig 3A, right vs left). First, both cortical and thalamic spindles were more spatially focal in the core system as only a small subset of neurons was involved in a typical spindle at any given time. In contrast, within the matrix system spindles were global (involving the entire cell population) and highly synchronous across all cell types. These results are consistent with our previous studies [5] and suggest that connectivity properties of thalamocortical projections determine the degree of synchronization in the cortical network. Second, spindle density was higher in the core system compared to the matrix system. At every spatial location in the cortical network of the core system, the characteristic time between spindles was shorter compared to that between spindles in the matrix system (Fig 3A left vs right). In order to quantify the spatial and temporal properties of spindles, we computed an estimated LFP as an average of the dendritic synaptic currents for every group of contiguous 100 cortical neurons. LFPs of the core system were estimated from the layer 3/4 cells’ dendrites and the LFP of the matrix system was computed from the dendrites of layer 5 neurons that were located in the superficial cortical layers (Fig 2). After applying a bandpass filter (6-15 Hz), the spatial properties of estimated core and matrix LFP (Fig 3C) closely matched the MEG and EEG recordings, respectively (Fig 1). In subsequent analyses, we used this estimated LFP to further examine the properties of the spindle oscillations in the core and matrix system.

### Spindle occurrence can be captured by a non-periodic process

We identified spindles in the estimated LFP using an automated spindle detection algorithm similar to those used in experimental studies (details are provided in the method section). The spindle density, as defined by the number of spindles occurring per minute of simulation time, was greater in the core compared to the matrix (Fig 4A) confirmed by a independent sample t-test (t(18) = 7.06, p<0.001 for across estimated LFP channels and t(2060)= 19.2, p<0.001 across all spindles). This pattern resembles the experimental observation that EEG spindles occur more frequently than MEG spindles. While the average spindle density was significantly different between the core and matrix, in both systems the distribution of inter-spindle intervals peaks was below 4s and has a long tail (Fig 4B). A two sample KS test comparing the distributions of inter-spindle intervals confirmed that the intervals were derived from different distributions (D_1128,932_=0.427, p<0.001). The peak ISI of the core system was shorter than then that of the matrix system, suggesting that the core experiences shorter and frequent quiescence periods than the matrix. Furthermore, maximum-likelihood fits of the probability distributions (red line in Fig 4B) confirmed that the intervals of spindle occurrence cannot be described by a normal distribution. The long tails of the distributions suggest that a Poisson like process, as oppose to a periodic process, is responsible for spindle generation. This observation is consistent with previous experimental results [14, 22] and suggests that our computational model replicates an essential statistical property observed in *in vivo* experiments

**Figure 4:**
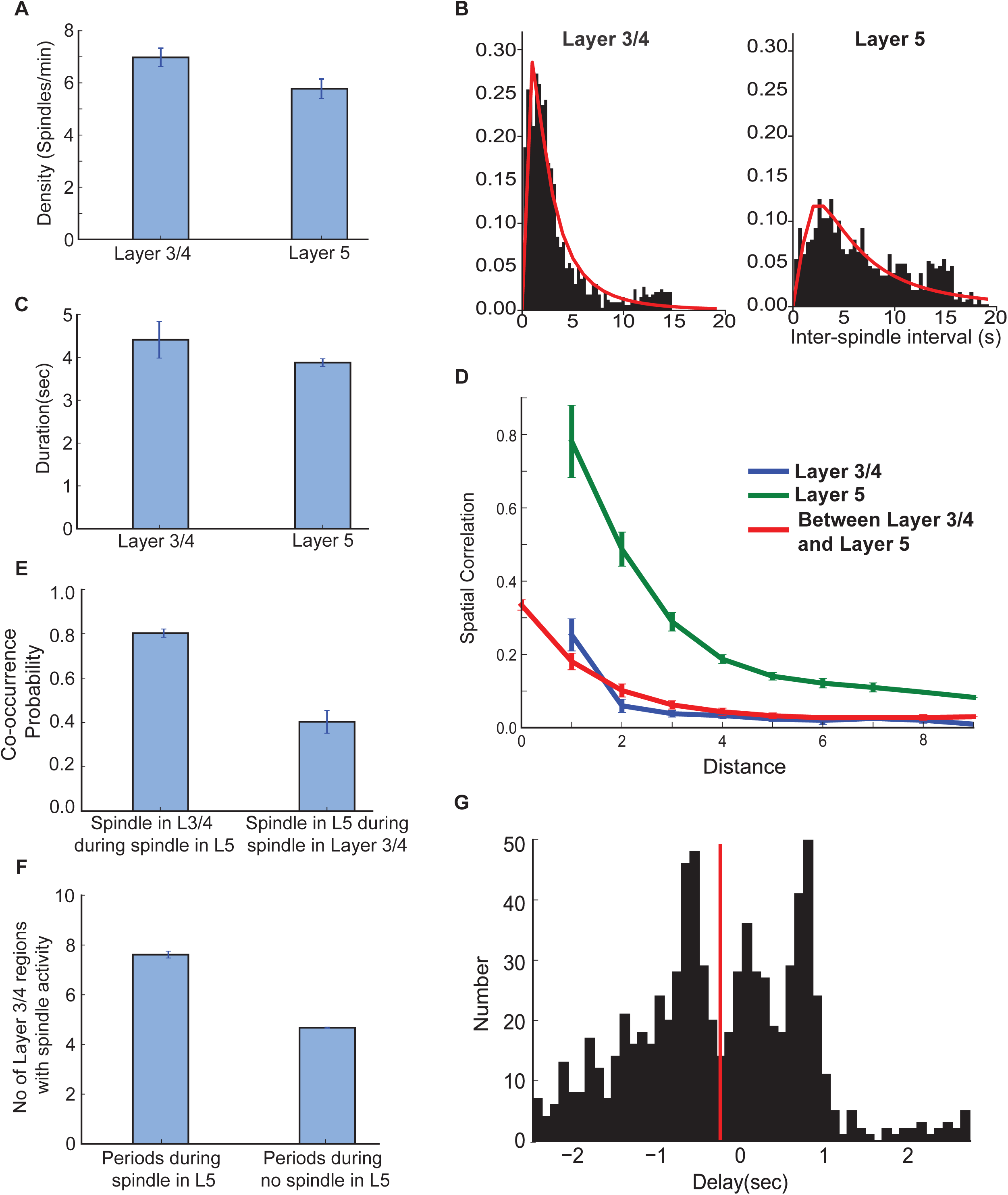
Statistical proprieties of sleep spindles in core and matrix systems. (A) Spindle density measured from the filtered averaged network activity in the core and matrix systems, with error bar indicating standard error across different network locations.(B)Histograms of the inter-spindle intervals show a non-normal distribution in the occurrence of spindles in both the core and matrix systems. (C) Spindle duration as measured by the onset to offset of spindle that was identified by a threshold measure (see method section for details) and was significantly different (). (D) Spatial correlation within and between layers measured as a function of distance between network sites. (F) Probability of spindle co-occurrences in the core and matrix systems. (G) Delay between the onset of spindles in the core and matrix, negative values indicate that the spindle in the core precedes that in the matrix.

Several other features of core and matrix spindles in the model simulations were similar to experimental recordings. The average spindle duration was significantly higher in the core compared to the matrix system (Fig 4C) confirmed by independent t-test (t(2060)=16.3, p<0.001). To quantify the difference in the spatial synchrony of spindles between core and matrix systems, we computed the spatial correlation [23] between the LFP groups at different distances (measured by the location of a neuron group in the network). The correlation coefficient diminished with distance for both systems (Fig 4D). However, the spindles in the core system were less spatially correlated overall compared to spindles in the matrix system.

### Co-occurring core and matrix spindles recruits large cortical region in the core system through recurrent excitation between layers

Simultaneous EEG and MEG measurements have found that about 50% of the MEG spindles co-occur with EEG, while about 85% of EEG spindles co-occur with MEG [24]. Further, spindle detected in EEG is observed to co-occur in about 66% more MEG channels than a spindle detected in MEG. Our model produces spindling patterns consistent with these features. The co-occurrence probability revealed that during periods of spindles in the matrix system, there was about 80% probability that core was also involved in spindles (Fig 4E). In contrast, there was only a 40% probability of observing a matrix spindle during a spindle in the core system. Independent t-test confirmed this difference between the systems across estimated LFP channels (t(14)=31.4, p<0.001). Further, we observed that the number of LFP channels that were activated during a spindle event in the core system was higher when spindles co-occurred in the matrix versus times when the spindles occurred only in the core (Fig 4F, t(14)=67.2, p<0.001). This suggests that co-occurrences of spindles in both systems are rare events but lead to the wide spread activation in both core and matrix when they occur.

Finally, we examined the delay between spindles in the core and matrix systems. We observed that on average (red line in Fig 4G), the spindle originated from the core system then spread to the matrix system with a mean delay of about 300 ms (delay was measured as the difference in onset times between co-occurring spindles within a window of 2,500 ms; negative delay values indicate spindles in which the core preceded matrix). The peak at -750 ms corresponds to spindles originating from core system, while the peak at 500 ms suggests that at some network sites, spindles originated in the matrix system and then spread to the core system.

In sum, these results suggest that spindles are more frequently initiated locally in the core system, then propagate from the core system to the matrix system which leads to the spindles spreading throughout the matrix system. Eventually, even regions in the core system that were not previously involved are recruited. These findings explain the experimental observation that spindles are observed in more MEG channels when they also present in EEG [24].

### Impact of thalamocortical and corticothalamic connection on spindle occurrence

We leveraged our model to examine factors that may influence spindle occurrences across cortical layers. The main difference between the core and matrix systems in the model was the breadth or fanout of the thalamic projections to the cortical network. Neuroanatomical studies suggest that the core system has focused projections while matrix system projects widely [12]. Here, we assessed the impacts of this characteristic by systematically varying the connection footprint of the thalamic matrix to superficial cortical regions, while holding the fanout of the thalamic core to layer 3/4 constant. We also modulated the corticothalamic projections in proportion to the thalamocortical projections. Using the estimated LFP from the cortical layers corresponding to core and matrix system, respectively, we quantified various spindle properties as the fanout was varied.

Spindle density (number of spindles per minute) in both layers was sensitive to matrix system fanout. ANOVA confirmed significant effects of the fanout and layer location, as well as an interaction between layer and fanout (fanout: F(6,112)= 66.4; p<0.01, Layer: F(1,112)= 65.18; p<0.01 and interaction F(6,112)= 22.8; p<0.01). When matrix and core thalamus have similar fanouts (ratio 1 and 2.5 in Fig 5B), we observed a slightly higher density of spindles in the matrix than in the core system. This observation is consistent with the differing circuits (see Fig 2), wherein the matrix system contains direct reciprocal projections connecting cortical and thalamic subpopulations and the core system routes indirect projections from cortical (layer III/IV) neurons through layer VI to the thalamic nucleus. When the thalamocortical fanout of the matrix system was increased to above ∼5 times the size of the core system, the density of spindles in the matrix system was reduced to around 4 spindles per minute. Interestingly, the density of spindles in the core system was also reduced when the thalamocortical fanout of the matrix system was further increased to above ∼10 times of that in the core system (ratio above 10 in Fig 5B). This suggests that spindle density in both systems is determined not only by the radius of thalamocortical vs. corticothalamic projections, but also by interactions between the systems among the cortical layers. We further expound on the role of these cortical connections in the next section.

**Figure 5:**
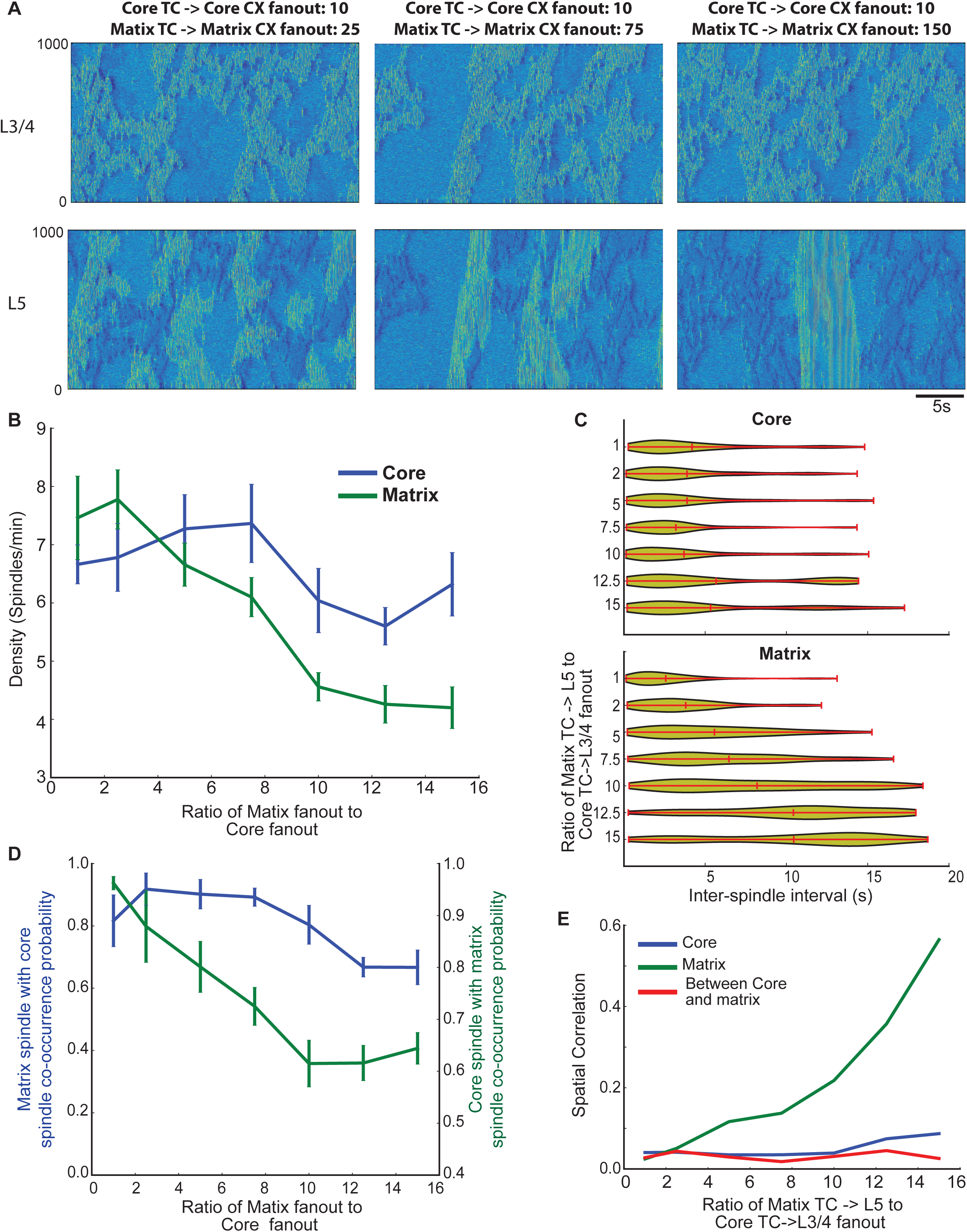
Effect of thalamocortical connectivity on the spindle properties. (A) Space-time plots of the network activity in the cortical layers for different fanout of the thalamocortical and corticothalamic connections. (B) Spindle density for different fanout conditions. The x-axis corresponds to the ratio of thalamic projection fanout in the matrix compared to the core system. The actual thalamocoritcal and corticothalamic radii for the different ratios (1,2.5, 5, 7.5, 10, 12.5, 15) were 10/2/10/2 (core TC->L3/4, L6->core TC, matrix TC->L5, L5->matrix TC), 10/2/25/5, 10/2/50/10, 10/2/75/15, 10/2/100/20, 10/2/125/25, 10/2/150/30. (C) Violin plots showing the distribution of inter-spindle intervals as a function of thalamocortical fanout. (D, E) Probability of spindle co-occurrence between core and matrix, and spatial correlation as a function of synaptic fanout.

We also examined effects of thalamocortical fanout on the distribution of inter-spindle intervals (Fig 5C). Although the mean value was largely independent on the projection radius, a long tailed distribution was observed for all values of fanout in the core. Contrastingly, in the matrix system the mean and the peak of the inter-spindle interval shifted to the right (longer intervals) with increases in fanout. With large fanouts, the majority of matrix system spindles had very long periods of silence (10-15s) between them. This suggests that thalamocortical fanout determines the peak of the inter-spindle interval distribution, but does not alter the stochastic nature of spindle occurrence.

The size of the thalamocortical fanout also influenced the co-occurrence of spindles in the core and matrix systems (Fig 5D). Increasing the fanout of the matrix system reduced spindle co-occurrence between two systems. This reduction resulted mainly from a lower spindle density in both layers. However, the co-occurrence of core spindles during matrix spindles was higher for all values of the fanout when matrix thalamocortical projections were at least 5 times broader than core projections. This suggests that the difference in spindle co-occurrence between EEG and MEG as observed in experiments [11] depends mainly on the difference in the radius of thalamocortical projections between core and matrix systems, while overall level of co-occurrence is determined by the interaction between cortical layers.

We examined how spatial correlations during periods of spindles vary depending on the fanout of thalamic projections. The spatial correlation measures the degree of synchronization in the estimated LFP between network locations as a function of the distance between locations. As expected, increasing distance reduced the spatial correlation (Fig 4D). We next measured the mean value of the spatial correlation for each fanout condition. The mean correlation increased as a function of fanout in the matrix system (Fig 5A). However, the spatial correlation within the core and between the core and matrix systems did not change with increases in fanout, suggesting that the spatial synchronization of core spindles is largely influenced by thalamocortical fanout but not by interactions between core and matrix systems as was observed for spindle density.

### Feedforward input to matrix critically determines spindles density

Does intra-cortical excitatory connectivity between layer 3/4 of the core system and layer 5 of the matrix system affect spindle occurrence? To answer this question, we first, varied the strength of excitatory connections (AMPA and NMDA) from core to matrix pyramidal neurons (Fig 6A-B). Here the reference point (or 100%) corresponds to the strength used in previous simulations, i.e. half the strength of a within-layer connection. The spindle density varied with the strength of the interlaminar connections (Fig 6A). For low connectivity strengths (below 100%), the spindle density of the matrix system was reduced significantly, while at high strengths (above 140%) the spindle density of the matrix system exceeded that of the control (100%). There were significant effects of connection strength, layer, and interaction on spindle density (connection strength: F(5,96)=24.7; p<0.01, layer: F(5,96)= 386.6; p<0.01 and interaction F(5,96)= 36.9; p<0.01) that suggest a layer-specific effect of modulating excitatory interlaminar connection strength. Similar to spindle density, spindle co-occurrence between core and matrix systems also increased as a function of the interlaminar connection strength, reaching 80% for both core and matrix at 150% connectivity. In contrast, changing the strength of excitatory connections from layer 5 to layer 3/4 had little effect on the spindle density, (Fig 6C). Taken together, these results suggest that the strength of cortical core-to-matrix excitatory connections is one of the critical factors determining spindle density and co-occurrence among spindles across both cortical lamina and the core/matrix systems.

**Figure 6:**
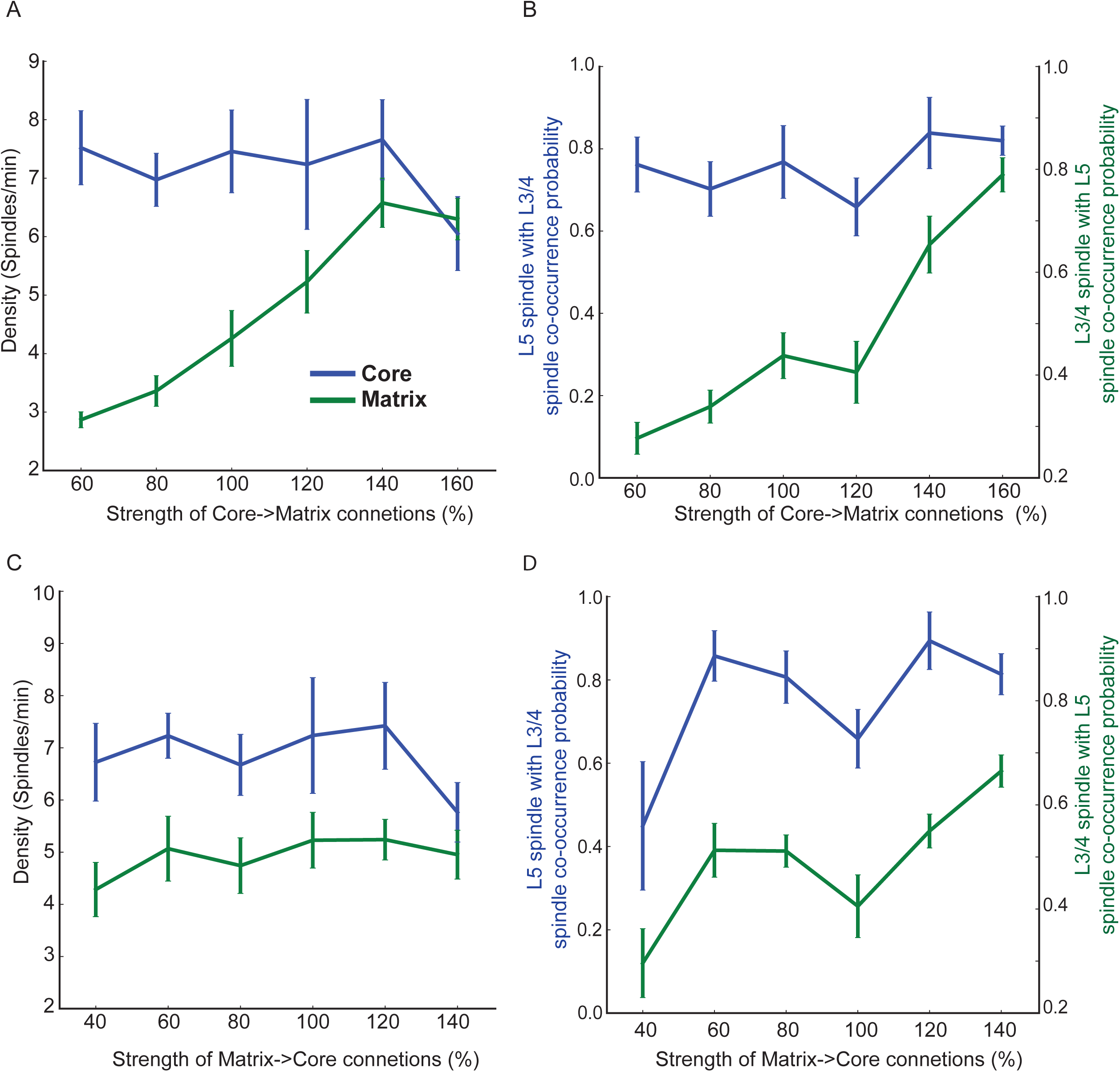
Effect of interlaminar connectivity on the spindle properties. (A,B) Spindle density and probability of co-occurrence between core and matrix as a function of strength of the AMPA and NMDA connections from L3/4 to L5. 100% interlaminar connectivity corresponds to half of the intralaminar connection strength (within L3/4 and L5). (C, D) Spindle density and probability of co-occurrence as a function of the AMPA and NMDA connection strength from L5 to L3/4 neurons.

## Discussion

Using EEG/MEG recordings in humans and computational modeling we demonstrated that properties of sleep spindle occurrence vary across cortical layers and are influenced by thalamocortical, corticothalamic and cortico-laminar connections. This study was motivated by empirical findings demonstrating that spindles measured in EEG and MEG have a different synchronization properties across cortical populations [11, 24]. EEG spindles occur less frequently and more synchronously in comparison to MEG. Our new study predicts that anatomical differences between the matrix thalamocortical system, which has broader projections that target the cortex superficially, and the core system, which consists of the focal projections that target the middle layers, can account for the differences between EEG and MEG signals. Furthermore, we discovered that the strength of cortico-cortico feedforward excitatory connections from the core to the matrix neurons determined spindle density in the matrix system, which may provide greater context to EEG recordings.

The origin of sleep spindle oscillatory activity has been linked to thalamic oscillators based on the broad range of experimental studies [2, 25, 26]. The excitatory and inhibitory connections between thalamic relay and reticular neurons are critical in generating the spindle oscillation [15, 18, 27, 28]. However, in intact brain, sleep spindles are critically shaped by the cortical networks. Indeed, the onset of a spindle oscillation and its termination are both dependent on the cortical input to the thalamus [3, 29]. Spindle oscillations in the thalamus are initiated when sufficiently strong activity in the cortex activates the thalamic network, and spindle termination is partially mediated by the desynchronization of thalamic input toward the end of spindle. In addition, thalamocortical interactions are known to be integral to the synchronization of spindles [5, 28]. In our new study, the core thalamocortical system had relatively higher spindle density produced by focal and strong thalamocortical and corticothalamic connections. Such a pattern of connectivity between core thalamus and middle cortical layers allowed input from a small region of the cortex to initiate and maintain focal spindles in the core system. In contrast, the matrix system had relatively weak and broad thalamocortical connections requiring synchronized activity in broader cortical regions in order to initiate spindle activity in the thalamus.

We previously reported [5] that (1) *within* a single spindle event the synchrony of the neuronal firing is higher in the matrix than in the core subsystem; (2) spindle are initiated in the core and with some delay in the matrix subsystem. The overal density of the core and matrix spindle events was, however, the same in this earlier model. In the new study we extended these previous results by explaining the difference in the global spatio-temporal structure of spindle activity between core and matrix. Thus, our new model predicts that the focal nature of the core thalamocortical connectivity can explain more frequent occurrence of spindles in the core than in the matrix susbsystem as observed in vivo. The strength of the core-to-matrix intracortical connections determined probability of the core spindles to “propagate” to the matrix subsystem. Futhermore, in our new model core spindles remained localized and have never involved the entire network, again in agreement with in vivo data.

We observed that the distribution of the inter-spindle intervals reflects a non-periodic stochastic process such as a Poisson process which is consistent with the previous experimental studies [14, 22]. The state of the thalamocortical network, as determined by the levels of the intrinsic and synaptic conductances, contributed to the stochastic nature of spindle occurrence. Building off our previous work [16], we chose the intrinsic and synaptic properties in the model that match those in stage 2 sleep, a brain state when concentrations of neuromodulating acetylcholine and monoamines are reduced [30-32]. As a consequence, the K-leak currents and excitatory intracortical connections were set higher compared to an awake-like state due to the reduction of acetylcholine and nor-epinephrine [33]. The high K-leak currents resulted in sparse spontaneous cortical activity during periods between spindles with occasional surges of local synchrony sustained by increased recurrent excitation within the cortex that occasionally induce spindle oscillations in the thalamus. Furthermore, the release of miniature EPSPs and IPSPs in the cortex was implemented as a Poission process that contributed to the stochastic nature of baseline activity. All these factors led to the variable inter-spindle interval with long periods of silence when activity in cortex was not sufficient to induce spindles. While it is known that excitable medium with noise has Poisson event distribution in reduced systems [34], we describe here that a detailed biophysical model of spindle generation lead to Poission process due to specific intrinsic and network properties.

Simultaneous EEG and MEG recordings as reported here and in the previous publications [11, 24] have revealed that the MEG spindles occur earlier compared to the EEG spindles and EEG spindles are seen in a higher number of MEG sensors compared to the spindles occurring only in the MEG data. This resembles our current findings, in which the number of regions that were spindling in the core system during a matrix spindle was higher than when there was no spindle in the matrix system. Further, the distribution of delays between the onset of spindles between the systems indicate that during matrix spindles some cortical neurons of the core system fired early, and presumably contributed to the initiation of the matrix spindle, while others fired late and were recruited. Taken together, all the evidence suggests a characteristic and complex spatiotemporal evolution of spindle activity during co-occurring spindles, where spindles in the core spread to the matrix that in turn activate wide regions in the core leading to synchronized activation across cortical layers that is reflected in the strong activity in both EEG and MEG. Thus, model predicts that co-occurring spindles could lead to recruitment of the large cortical areas, which indeed has been reported in previous studies [23, 35]. At the same time, local spindles occurring in the model within deep layer may correspond to the local spindles observed in some studies [36], or even hidden from empirical recordings because of their localized and low amplitude properties. Finally, the regional differences in thalamocorical and cortical connections could explain characteristic regional and spatial patterns of spindles observed in human recordings [37].

Increase in spindle density following learning is a robust experimental finding that demonstrates the role of sleep in memory consolidation [2, 6, 7, 10, 38, 39]. However, the neural mechanisms that increase in spindle density after learning are not known. Projections of the hippocampal CA1 region have efferents to both superficial and deep layers of the prefrontal cortex [40, 41]. In addition, experiments with simultaneous recordings from cortex and hippocampus report that during NREM sleep, the two structures show coordinated activity [42, 43] that underpins spike sequence replay and the reactivation of memories that were learned when awake [44]. In our study, we found that activation of the layer 3/4 of the neocortex triggers spindles that propagate between core and matrix systems and eventually lead to spindle recruitment in wide regions of the neocortex. Based on these findings, we predict that hippocampal input to the superficial cortical layers (layer 2/3/4) during NREM sleep can induce local activation and spindles in the core system, which then propagate to the matrix system thereby activating large cortical regions in both layers, potentially contribution to memory consolidation.

Elevated spindle density may arise due to the changes in the cortical microcircuit, of which excitatory interlaminar connections form the main component [45]. This circuit is implicated as a site of the sensory coding and learning [46]. In this study, we identified that connection strength from the core to the matrix, but not from the matrix to the core, was critical in determining spindle density. This predicts that the increase in spindle density following a hippocampal-dependent task may arise from the strengthening of feedforward projections connecting superficial layers to the cortical matrix.

In sum, our study identified a rich set of the local and global network mechanisms involved in spindle propagation and the interactions among spindles in cortical structures. While spindle activity in the model arises from the thalamic circuits, our study supports the idea that thalamocortical and intracortical projections significantly shape the properties of spindling activity and that this may explain the characteristic changes of spindles density associated with sleep-related memory replay.

## Materials and Methods

### Computational models

*Intrinsic currents* – thalamus. Conductance based models of thalamocortical neuron (TC) and thalamic reticular neuron (RE) including one compartment are described by the equation:

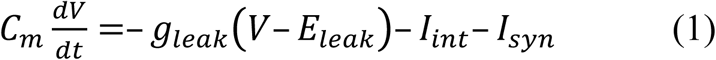

where the membrane capacitance, C_m_, is equal to 1 μF/cm^2^, non-specific (mixed Na^+^ and Cl^-^) leakage conductance, g_leak_, is equal to 0.01 mS/cm^2^ for TC cells and 0.05 mS/cm^2^ for RE cells, and the reversal potential, E_leak_, is equal to – 70 mV for TC cells and – 77 mV for RE cells. I_int_ is the sum of active intrinsic currents, and I_syn_ is the sum of synaptic currents. The area of a RE cell and a TC cell was 1.43×10-4 cm^2^ and 2.9×10-4 cm^2^, respectively [21, 47].

Both RE and TC cells include fast sodium current, I_Na_, a fast potassium current, I_K_, a low-threshold Ca^2+^ current I_T_, and a potassium leak current, I_KL_ = g_KL_ (V – E_KL_), where E_KL_ = – 95 mV. In addition, a hyperpolarization-activated cation current, I_h_, was also included in TC cells. For TC cells, the maximal conductances are g_K_ = 10 mS/cm^2^,g_Na_= 90 mS/cm^2^, g_T_= 2.2 mS/cm^2^, g_h_ = 0.017 mS/cm^2^, g_KL_ = 0.03 mS/cm^2^. For RE cells, the maximal conductances are g_K_ = 10 mS/cm^2^, g_Na_ = 100 mS/cm^2^, g_T_ = 2.3 mS/cm^2^, g_leak_= 0.005 mS/cm^2^.The expressions of voltage-and Ca^2+^-dependent transition rates for all currents are given in [21, 47].

*Intrinsic currents* – cortex. The cortical pyramidal cells (PY) and interneurons (IN) were represented by two-compartment model with channels simulated by Hodgkin–Huxley kinetics. Each compartment was described by Eq (1) with an additional current from adjacent compartment, I_d,s_ = g_c_(V_d,s_– V_s,d_), where V_d_ (resp. V_s_) is the voltage of the dendritic (resp. axo-somatic) compartment. The axo-somatic and dendritic compartments were coupled by an axial current with conductance g_c_.

The PY neurons and INs contained the fast Na^+^ channels, I_Na_, with a higher density in the axosomatic compartment than in dendritic compartment. In addition, a fast delayed rectifier potassium K^+^ current, I_K_, was present in the axosomatic compartment. A persistent sodium current, I_Na(p)_, was included in both axosomatic and dendritic compartments. A slow voltage-dependent non-inactivating K^+^ current, I_Km_, a slow Ca^2+-^ dependent K^+^ current, I_Kca_, a high-threshold Ca^2+^ current, I_HVA_, and a potassium leak current, I_KL_ = g_KL_(V-E_KL_) were included in the dendritic compartment only. The expressions of the voltage- and Ca^2+^-dependent transition rates for all currents are given in Timofeev et al. [48]. For axosomatic compartment, the maximal conductances and passive properties were S_soma_ = 1.0×10 -6 cm^2^, g_Na_ = 3000 mS/cm^2^, g_K_ = 200 mS/cm^2^, g_Na(p)_ = 0.07 mS/cm^2^. For dendritic compartment: Cm=0.75 μF/cm^2^, gL = 0.033 mS/cm^2^, gKL = 0.0025 mS/cm^2^, S_dend_ = S_soma_ × R, g_HVA_ = 0.01 mS/cm^2^, g_Na_ = 1.5 mS/ cm^2^, g_Kca_ =0.3 mS/cm^2^, g_Km_ = 0.01 mS/cm^2^, g_Na(p)_ = 0.07 mS/ cm^2^, E_KL_ = – 95 mV. E_leak_ was – 68 mV for PYs and – 75 mV for INs [21, 47]. For interneurons, no I_Na(p)_ was included. The resistance (r) between compartments was set to 10 MΩ.

The firing properties of this model depended on the coupling conductance between compartments (g_c_ = 1/r) and the ratio of dendritic area to the axosomatic area R (Mainen and Sejnowski, 1996). We used a model of a regular-spiking neuron for PY cells (r = 165) and a model of a fast spiking neuron for IN cells (r = 50).

Synaptic currents. All synaptic currents were calculated according to:

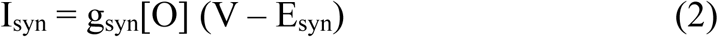

where g_syn_ is the maximal conductance, [O] is the fraction of open channels, and E_syn_ is the reversal potential. In RE and PY cells, reversal potential was 0 mV for AMPA and NMDA receptors, and –70 mV for GABAA receptors. For TC cells, the reversal potential was –80 mV for GABAA receptors, and –95 mV for GABAB receptors. A simple phenomenological model characterizing short-term depression of intracortical excitatory connections was also included in the model. According to this, a maximal synaptic conductance was multiplied to a depression variable, D, which represents the amount of available synaptic resources. Here, D = 1 – (1 – D_i_ (1 – U)) exp (– (t – t_i_)/τ) where U =0.2 was the fraction of resources used per action potential, τ = 500 msec was the time constant of recovery of the synaptic resources, D_i_ was the value of D immediately before the ith event, and (t – t_i_) is the time after i^th^ event.

GABAA, NMDA, and AMPA synaptic currents were modeled by the first-order activation schemes. Dependence of postsynaptic voltage for NMDA receptors was 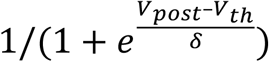, where V_th_ = – 25 mV, δ = 12.5 mV. GABAB receptors were modeled by a higher-order reaction scheme that considers the activation of K^+^ channels by G-proteins. The equations for all synaptic currents were given in [48].

Spontaneous miniature EPSPs and IPSPs were also set in the current model. The arrival times of miniature EPSPs and IPSPs followed the Poisson process (Stevens, 1993), with time-dependent mean rate μ=(2/(1+exp(-(t-t_0_)/400))-1)/250 where t was real time and t_0_ was timing of the last presynaptic spike occurring.

*Network geometry*. The thalamocortical network model was constructed using several layers of one-dimensional networks that included cortical pyramidal cells (layer II-IV; layer V; layer VI), cortical interneurons (layer II-IV; layer V; layer VI), thalamocortical neurons (with matrix and core systems) and thalamic reticular neurons. A general connecting scheme among different populations of neurons is introduced in Fig. 5. In total, there were 3000 pyramidal neurons (1000 in each population), 400 interneurons (200 in each population), 400 thalamic reticular neurons, and 400 thalamocortical neurons (200 in each system).

The maximal conductances were (subscript indicates cell types and superscript indicates type of connection): connections within cortical layers had following conductances 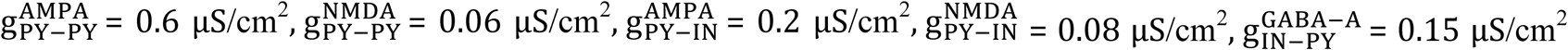; conductances for connections between layers were: 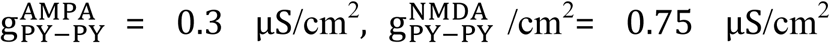. Thalamocortical and corticothalamic connections for core and matrix system had following conductances: 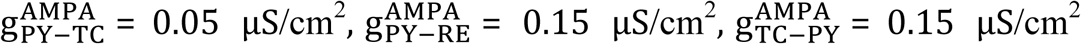. Connections within thalamus for both core and matrix system had following conductances: 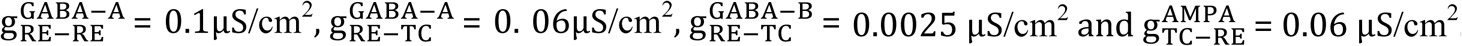 and 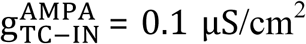.

## Experimental Methods

Extracranial electromagnetic fields were recorded in 4 healthy adults (3 female). Subjects did not report any neurological problems including sleep disorders, epilepsy, or substance dependence. In addition, subjects did not consume caffeine or alcohol on the day of recording. Written informed consent approved by the human research committee at Partners Healthcare Network was obtained from all subjects before participation. A whole-head MEG system with integrated EEG cap (Elekta Neuromag) was used to collect 204 planar gradiometers and 60 EEG channels. EEG data were referenced to an averaged mastoid. Additional details concerning data collection can be found in [49]. Sleep staging was performed by three neurologists according to standard criteria (Rechtschaffen and Kales, 1968). Data analyzed came from a 17.5 ± 3.4 (mean ± SD) minute period of stage 2 sleep.

Data were acquired at 603.107 Hz. Gross artifacts and bad channels were excluded manually. The continuous data were band-pass filtered to between 0.1 and 30 Hz and ICA (Delorme and Makeig, 2004) was used to remove the ECG component. Spindles were automatically detected in each MEG and EEG channel using a method modified from [36].The 10-16 Hz analytic signal was extracted from the data using the Hilbert transform and the complex envelope obtained by computing its modulus. The spindle band envelope was smoothed with a Gaussian kernel (300 ms width, 40 ms σ). Putative spindles were initially marked as contiguous regions of the smoothed spindle-band envelope where the envelope amplitude was more than 2 standard deviations above the mean. Marked regions were then expanded until amplitude dropped below 1 standard deviation above the mean. Putative spindles shorter than 500 ms and longer than 2 s were excluded from further analysis. Inter-spindle intervals (ISIs) were computed from spindle center to center. Outlying ISIs longer than 20 seconds were excluded, as these are likely caused by false negatives in spindle detection. ISIs from all subjects and all channels were pooled together to form a single distribution for EEG and gradiometer data, respectively.

## ACKNOWLEDGMENTS

This work was supported by grants from ONR (MURI: N000141310672) and NIH (MH099645 and EB009282).

## References

1. Timofeev I, Steriade M. Low-frequency rhythms in the thalamus of intact-cortex and decorticated cats. J Neurophysiol. 1996;76(6):4152–68.

2. Kim U, Bal T, McCormick DA. Spindle waves are propagating synchronized oscillations in the ferret LGNd in vitro. J Neurophysiol. 1995;74(3):1301–23. PubMed PMID: 7500152.

3. Bonjean M, Baker T, Lemieux M, Timofeev I, Sejnowski T, Bazhenov M. Corticothalamic feedback controls sleep spindle duration in vivo. J Neurosci. 2011;31(25):9124–34. Epub 2011/06/24. doi: 10.1523/JNEUROSCI.0077-11.2011. PubMed PMID: 21697364.

4. Contreras D, Destexhe A, Sejnowski TJ, Steriade M. Spatiotemporal patterns of spindle oscillations in cortex and thalamus. J Neurosci. 1997;17(3):1179–96. PubMed PMID: 8994070.

5. Bonjean M, Baker T, Bazhenov M, Cash S, Halgren E, Sejnowski T. Interactions between core and matrix thalamocortical projections in human sleep spindle synchronization. J Neurosci. 2012;32(15):5250–63. doi: 10.1523/JNEUROSCI.6141-11.2012. PubMed PMID: 22496571; PubMed Central PMCID: PMCPMC3342310.

6. Eschenko O, Molle M, Born J, Sara SJ. Elevated sleep spindle density after learning or after retrieval in rats. J Neurosci. 2006;26(50):12914–20. doi: 10.1523/JNEUROSCI.3175-06.2006. PubMed PMID: 17167082.

7. Fogel SM, Smith CT. Learning-dependent changes in sleep spindles and stage 2 sleep. Journal of sleep research. 2006;15(3):250–5.

8. Clemens Z, Fabo D, Halasz P. Overnight verbal memory retention correlates with the number of sleep spindles. Neuroscience. 2005;132(2):529–35. doi: 10.1016/j.neuroscience.2005.01.011. PubMed PMID: 15802203.

9. Gais S, Molle M, Helms K, Born J. Learning-dependent increases in sleep spindle density. J Neurosci. 2002;22(15):6830–4. doi: 20026697. PubMed PMID: 12151563.

10. Mednick SC, McDevitt EA, Walsh JK, Wamsley E, Paulus M, Kanady JC, et al. The critical role of sleep spindles in hippocampal-dependent memory: a pharmacology study. J Neurosci. 2013;33(10):4494–504. doi: 10.1523/JNEUROSCI.3127-12.2013. PubMed PMID: 23467365; PubMed Central PMCID: PMCPMC3744388.

11. Dehghani N, Cash SS, Rossetti AO, Chen CC, Halgren E. Magnetoencephalography demonstrates multiple asynchronous generators during human sleep spindles. Journal of neurophysiology. 2010;104(1):179–88.

12. Jones EG. The thalamic matrix and thalamocortical synchrony. Trends Neurosci. 2001;24(10):595–601. PubMed PMID: 11576674.

13. Cash SS, Halgren E, Dehghani N, Rossetti AO, Thesen T, Wang C, et al. The human K-complex represents an isolated cortical down-state. Science. 2009;324(5930):1084–7. doi: 10.1126/science.1169626. PubMed PMID: 19461004; PubMed Central PMCID: PMCPMC3715654.

14. Panas D, Malinowska U, Piotrowski T, Zygierewicz J, Suffczynski P. Statistical analysis of sleep spindle occurrences. PLoS One. 2013;8(4):e59318. doi: 10.1371/journal.pone.0059318. PubMed PMID: 23560045; PubMed Central PMCID: PMCPMC3613364.

15. Bazhenov M, Timofeev I, Steriade M, Sejnowski T. Spiking-bursting activity in the thalamic reticular nucleus initiates sequences of spindle oscillations in thalamic networks. Journal of neurophysiology. 2000;84(2):1076–87.

16. Krishnan GP, Chauvette S, Shamie I, Soltani S, Timofeev I, Cash SS, et al. Cellular and neurochemical basis of sleep stages in the thalamocortical network. Elife. 2016;5. doi: 10.7554/eLife.18607. PubMed PMID: 27849520; PubMed Central PMCID: PMCPMC5111887.

17. Steriade M, McCormick DA, Sejnowski TJ. Thalamocortical oscillations in the sleeping and aroused brain. Science. 1993;262(5134):679–85.

18. Destexhe A, Bal T, McCormick DA, Sejnowski TJ. Ionic mechanisms underlying synchronized oscillations and propagating waves in a model of ferret thalamic slices. J Neurophysiol. 1996;76(3):2049–70. PubMed PMID: 8890314.

19. Destexhe A, Contreras D, Steriade M. Mechanisms underlying the synchronizing action of corticothalamic feedback through inhibition of thalamic relay cells J Neurophysiol. 1998;79(2):999–1016.

20. Destexhe A, Mainen ZF, Sejnowski TJ. Synthesis of models for excitable membranes, synaptic transmission and neuromodulation using a common kinetic formalism. J Comput Neurosci. 1994;1(3):195–230. PubMed PMID: 8792231.

21. Bazhenov M, Timofeev I, Steriade M, Sejnowski TJ. Model of thalamocortical slow-wave sleep oscillations and transitions to activated States. J Neurosci. 2002;22(19):8691–704. PubMed PMID: 12351744.

22. Himanen SL, Virkkala J, Huupponen E, Niemi J, Hasan J. Occurrence of periodic sleep spindles within and across non-REM sleep episodes. Neuropsychobiology. 2003;48(4):209–16. doi: 74639. PubMed PMID: 14673219.

23. Muller L, Piantoni G, Koller D, Cash SS, Halgren E, Sejnowski TJ. Rotating waves during human sleep spindles organize global patterns of activity that repeat precisely through the night. eLife. 2016;5:e17267.

24. Dehghani N, Cash SS, Halgren E. Emergence of synchronous EEG spindles from asynchronous MEG spindles. Hum Brain Mapp. 2011;32(12):2217–27. doi: 10.1002/hbm.21183. PubMed PMID: 21337472; PubMed Central PMCID: PMCPMC3798068.

25. Steriade M, Jones EG, Llinas R. Thalmic oscillations and signaling. New York: John Wiley & Sons; 1990. 431 p.

26. Timofeev I, Grenier F, Steriade M. Disfacilitation and active inhibition in the neocortex during the natural sleep-wake cycle: an intracellular study. Proc Natl Acad Sci U S A. 2001;98(4):1924–9. doi: 10.1073/pnas.041430398. PubMed PMID: 11172052; PubMed Central PMCID: PMCPMC29358.

27. McCormick DA, Bal T. Sleep and arousal: thalamocortical mechanisms. Annual review of neuroscience. 1997;20(1):185–215.

28. Destexhe A, Contreras D, Sejnowski TJ, Steriade M. A model of spindle rhythmicity in the isolated thalamic reticular nucleus. Journal of neurophysiology. 1994;72(2):803–18.

29. Timofeev I, Bazhenov M, Sejnowski T, Steriade M. Contribution of intrinsic and synaptic factors in the desynchronization of thalamic oscillatory activity. Thalamus & related systems. 2001;1(1):53–69.

30. Szymusiak R, Alam N, McGinty D. Discharge patterns of neurons in cholinergic regions of the basal forebrain during waking and sleep. Behav Brain Res. 2000;115(2):171–82. PubMed PMID: 11000419.

31. Vazquez J, Baghdoyan HA. Basal forebrain acetylcholine release during REM sleep is significantly greater than during waking. Am J Physiol Regul Integr Comp Physiol. 2001;280(2):R598–601. PubMed PMID: 11208592.

32. Takahashi K, Lin JS, Sakai K. Neuronal activity of histaminergic tuberomammillary neurons during wake-sleep states in the mouse. J Neurosci. 2006;26(40):10292–8. doi: 10.1523/JNEUROSCI.2341-06.2006. PubMed PMID: 17021184.

33. McCormick DA. Neurotransmitter actions in the thalamus and cerebral cortex and their role in neuromodulation of thalamocortical activity. Progress in neurobiology. 1992;39(4):337–88.

34. Lindner B, Garcia-Ojalvo J, Neiman A, Schimansky-Geier L. Effects of noise in excitable systems. Physics reports. 2004;392(6):321–424.

35. Contreras D, Steriade M. Spindle oscillation in cats: the role of corticothalamic feedback in a thalamically generated rhythm. The Journal of physiology. 1996;490(Pt 1):159–79.

36. Andrillon T, Nir Y, Staba RJ, Ferrarelli F, Cirelli C, Tononi G, et al. Sleep spindles in humans: insights from intracranial EEG and unit recordings. Journal of Neuroscience. 2011;31(49):17821–34.

37. Piantoni G, Halgren E, Cash SS. The contribution of thalamocortical core and matrix pathways to sleep spindles. Neural plasticity. 2016;2016.

38. Cox R, Hofman WF, de Boer M, Talamini LM. Local sleep spindle modulations in relation to specific memory cues. Neuroimage. 2014;99:103–10.

39. Nishida M, Walker MP. Daytime naps, motor memory consolidation and regionally specific sleep spindles. PLoS one. 2007;2(4):e341.

40. Jay TM, Witter MP. Distribution of hippocampal CA1 and subicular efferents in the prefrontal cortex of the rat studied by means of anterograde transport of Phaseolus vulgaris-leucoagglutinin. Journal of Comparative Neurology. 1991;313(4):574–86.

41. Parent MA, Wang L, Su J, Netoff T, Yuan LL. Identification of the hippocampal input to medial prefrontal cortex in vitro. Cereb Cortex. 2010;20(2):393–403. doi: 10.1093/cercor/bhp108. PubMed PMID: 19515741; PubMed Central PMCID: PMCPMC2803736.

42. Siapas AG, Wilson MA. Coordinated interactions between hippocampal ripples and cortical spindles during slow-wave sleep. Neuron. 1998;21(5):1123–8.

43. Sirota A, Csicsvari J, Buhl D, Buzsáki G. Communication between neocortex and hippocampus during sleep in rodents. Proceedings of the National Academy of Sciences. 2003;100(4):2065–9.

44. Ji D, Wilson MA. Coordinated memory replay in the visual cortex and hippocampus during sleep. Nature neuroscience. 2007;10(1):100–7.

45. Thomson AM, Lamy C. Functional maps of neocortical local circuitry. Frontiers in neuroscience. 2007;1:2.

46. Harris KD, Mrsic-Flogel TD. Cortical connectivity and sensory coding. Nature. 2013;503(7474):51–8.

47. Chen J-Y, Chauvette S, Skorheim S, Timofeev I, Bazhenov M. Interneuron-mediated inhibition synchronizes neuronal activity during slow oscillation. The Journal of Physiology. 2012;590(16):3987–4010.

48. Timofeev I, Grenier F, Bazhenov M, Sejnowski TJ, Steriade M. Origin of slow cortical oscillations in deafferented cortical slabs. Cerebral cortex (New York, NY: 1991). 2000;10(12):1185–99.

49. Dehghani N, Cash SS, Halgren E. Topographical frequency dynamics within EEG and MEG sleep spindles. Clinical Neurophysiology. 2011;122(2):229–35.

